# Label-free differential imaging of cellular components in mouse brain tissue by wide-band photoacoustic microscopy

**DOI:** 10.1101/2023.02.27.530195

**Authors:** Yajing Liu, Terence T W Wong, Junhui Shi, Yun He, Liming Nie, Lihong V. Wang

## Abstract

Mapping diverse cellular components with high spatial resolution is important to interrogate biological systems and study disease pathogenesis. Conventional optical imaging techniques for mapping biomolecular profiles with differential staining and labeling methods are cumbersome. Different types of cellular components exhibit distinctive characteristic absorption spectra across a wide wavelength range. By virtue of this property, a lab-made wide-band optical-resolution photoacoustic microscopy (wbOR-PAM) system, which covers wavelengths from the ultraviolet and visible to the shortwave infrared regions, was designed and developed to capture multiple cellular components in 300-μm-thick brain slices at nine different wavelengths without repetitive staining and complicated processing. This wbOR-PAM system provides abundant spectral information. A reflective objective lens with an infinite conjugate design was applied to focus laser beams with different wavelengths, avoiding chromatic aberration. The molecular components of complex brain slices were probed without labeling. The findings of the present study demonstrated a distinctive absorption of phospholipids, a major component of the cell membrane, brain, and nervous system, at 1690 nm and revealed their precise distribution with microscopic resolution in a mouse brain, for the first time. This novel imaging modality provides a new opportunity to investigate important biomolecular components without either labeling or lengthy specimen processing, thus, laying the groundwork for revealing cellular mechanisms involved in disease pathogenesis.

## Introduction

The spatial distributions and interactions of major cellular constituents including lipids, proteins, and nucleic acids, play a crucial role in exploring the function and structure of tissues, and the pathological mechanisms of diseases in humans^1-3^. Aberrant spatial distributions of cellular components have been associated with alterations of the cellular system and with human diseases^4^. Thus, mapping elementary cellular components accurately is essential to obtain relevant information for diagnostic pathology. Histopathological evaluation remains the clinical gold standard for diagnosis of diverse pathologies such as breast cancer, brain glioma, and cerebral atherosclerosis^5, 6^. However, the lesions of unstained tissue sections cannot be observed under a conventional optical microscope. Many specific staining techniques have been developed to preferentially stain different cellular components. For example, hematoxylin and eosin (H&E) staining is a classical histological technique that is extensively used to recognize cellular nuclei^7^; lipids are identified by Oil red O^8^; luxol fast blue staining identifies myelin^9^.

However, tissue specimen preparation is a time-consuming and cumbersome process. Moreover, the binding of dyes to target molecules is complicated, involving physical and chemical interactions. In addition, tissue specimens are easily deformed and ruptured during the preparation, subjecting histopathological images to various degrees of inaccuracy. In paraffin-embedded slices, organic solvents such as alcohols can remove lipids and destroy antigens easily^10^. The frozen section technique is suitable for fresh tissue but more demanding and tedious^11^. Formation of ice crystals may cause damage to cellular structures^12^. Furthermore, staining methods to highlight all cellular components are not readily available or feasible. Thus far, most of the optical imaging techniques are unable to differentiate different cellular components with high specificity.

Photoacoustic (PA) imaging (PAI), which is based on the thermoelastic expansion of an object caused by the absorption of light, is a rapidly-evolving imaging modality for early-stage theranostics and clinical applications^13-16^. PAI is a hybrid, non-ionizing, and non-invasive molecular imaging technique with scalable spatial resolution at depths beyond reach by conventional high-resolution optical imaging. By using appropriate wavelengths based on distinctive molecular absorption, optical-resolution PA microscopy (OR-PAM) can provide high-resolution, high-sensitivity and high-contrast images of cellular components without labeling. By taking advantage of the endogenous contrasts, PAI has been used to image different biological components, including hemoglobin, melanin, DNA/RNA, and cytochromes, at multiple length scales ranging from organelles to tissues^17-20^.

Wide-band OR-PAM (wbOR-PAM) takes advantage of the specific absorption peak of endogenous contrasts and captures these cellular components under multi-wavelength laser irradiation with high spatial resolution^21^. Upon illumination, a biological tissue specimen produces strong PA signals at matched wavelengths^22^. The molecules showing stronger PA amplitudes under their respective wavelengths are then identified. Consequently, endogenous contrasts can be spectrally separated without repetitive staining.

Different molecules exhibit unique absorption fingerprints due to characteristic chemical bonds. Lipids, such as cholesterol, phospholipids, and triglycerides, are known to exhibit characteristic absorption spectra that are similar to each other; however, with distinctive peaks in the shortwave infrared (SWIR) region. Phospholipids are the major constituent of the cell membrane and central nervous system^23,24^. The majority of previous studies have recorded phospholipids under the general absorption spectral range of lipids from 1210 to 1720 nm^25-27^. Nevertheless, none or only a limited number of the studies investigated the unique absorption of phospholipids separately. Herein, for the first time, we designed label-free wbOR-PAM with wide-band wavelengths (210–2600 nm), which spans from ultraviolet (UV) through visible (VIS) to SWIR wavelengths, to elucidate the spatial distribution of cellular components of mouse brain slices at 9 distinct wavelengths without complicated sectioning and multiple staining. The results reveal the characteristic absorption of phospholipids at 1690 nm and showed the distribution in the brain, which was in good agreement with the results based on myelin staining.

For biomolecular imaging, the optical window for the region of interest ranges between 200 and 2000 nm based on the absorption of the targeted biomolecules. Our wbOR-PAM system enables simultaneous imaging and differentiation of multiple absorbers existing in the biological specimen. With multiple wavelengths, this novel imaging technique provides a promising strategy to unravel cellular components, which paves a pathway for identifying subtle alteration in these elementary components of life and recognizing novel pathogenic mechanisms of human brain diseases.

## Results

As illustrated in Fig. 1, the label-free wbOR-PAM system works in transmission mode. The biological specimen is illuminated with a pulsed laser beam from the bottom, generating acoustic waves, detected by the upper confocally-aligned ultrasonic transducer. The laser beam is focused through a reflective objective lens. In contrast to a transmissive optical lens, the reflective objective lens consists of two mirrors to focus a broadband laser beam to a diffraction-limited spot without chromatic aberration. To reflect the UV light efficiently, a parabolic mirror (MPD149-F01, Thorlabs, Inc., USA) with UV enhanced aluminum coating was used in the light path. A three-axis motorized stage is triggered by a control box (sbRIO-9623, National Instruments, Corp., USA) to guide laser scanning. One-dimensional PA signals (A-lines) and B-scan images are shown on the LabVIEW interface in real time. Using an optical parametric oscillator (OPO) laser (210–2600 nm), the wbOR-PAM system allows identification of various endogenous contrasts based on their characteristic absorption spectra.

**Fig. 1.**
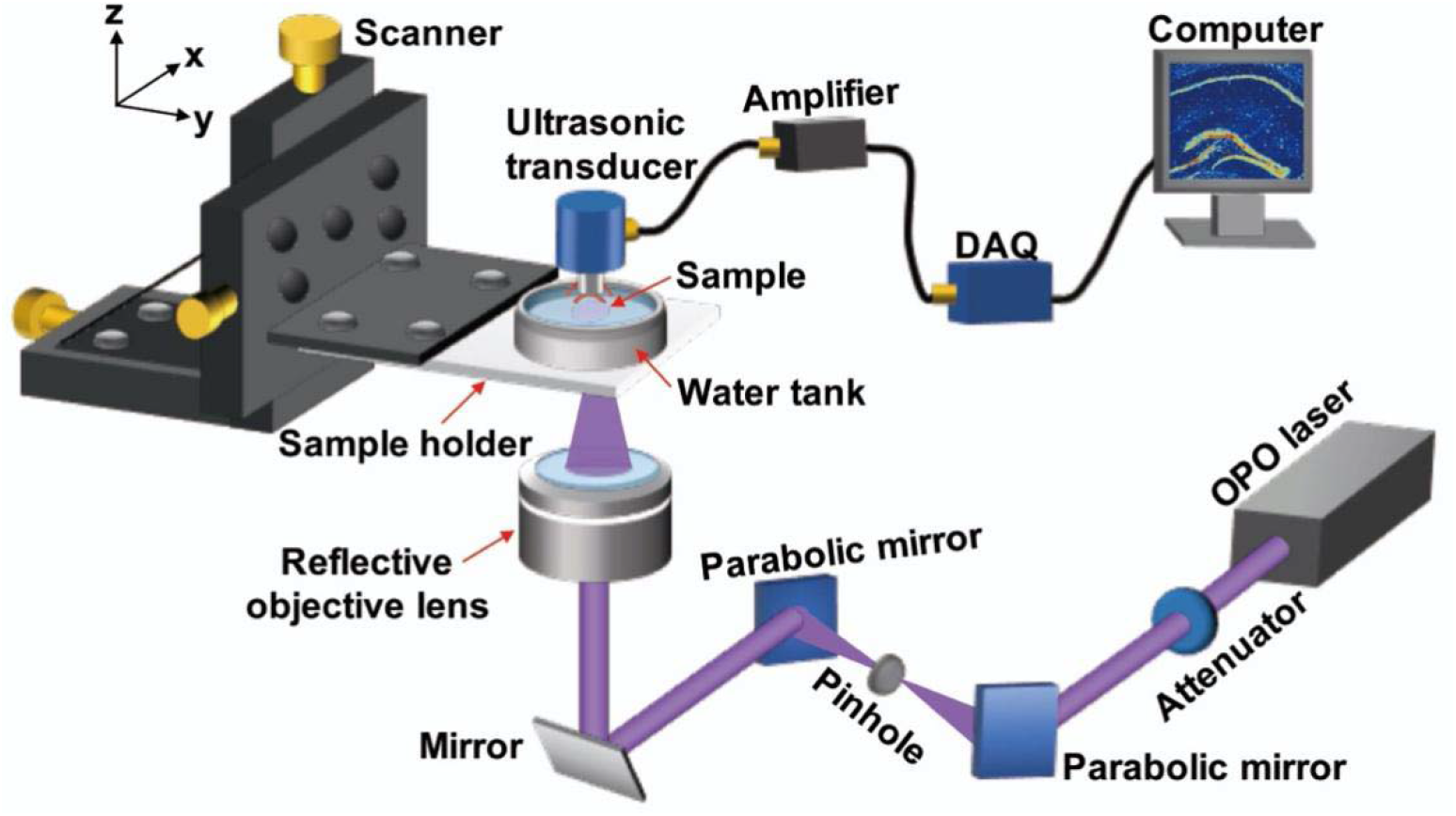
Schematic of the wide-band optical-resolution photoacoustic microscopy (wbOR-PAM) system with an OPO laser (210–2600 nm). DAQ: data acquisition card.

The wbOR-PAM images of paraffin-embedded 5-μm-thick mouse brain slices were acquired at the characteristic wavelengths (Fig. 2). The region of interest was 3.1 mm × 3.1 mm. The selected wavelengths covered the UV-VIS-near-infrared (NIR) regions (225, 266, 280, 420, 450, 1210 nm) to map the distribution of different cellular components (Fig. 2a-f). DNA and RNA exhibit strong absorption in the UV range (around 260 nm), highlighting cell nuclei^28^. Histopathologically, cancer is considerably associated with morphological and structural alterations of nuclei^29^. Due to the presence of aromatic amino acids, protein exhibits strong absorption in the UV region of 250–350 nm with maximum absorption at 280 nm^30^. Cytochrome, which is involved in the critical metabolism and biotransformation of a wide range of endogenous and exogenous substances^31^, predominantly absorbs at 420–450 nm. Lipids, which show its potential application in atherosclerosis diagnosis^26^, exhibit strong absorption in the NIR region (1210 nm). At their characteristic wavelengths, DNA/RNA, protein, cytochrome, and lipids can be captured with a high spatial resolution. These results indicate the high performance of wbOR-PAM, which is capable of providing more detailed cellular information compared with conventional single-wavelength OR-PAM.

**Fig. 2.**
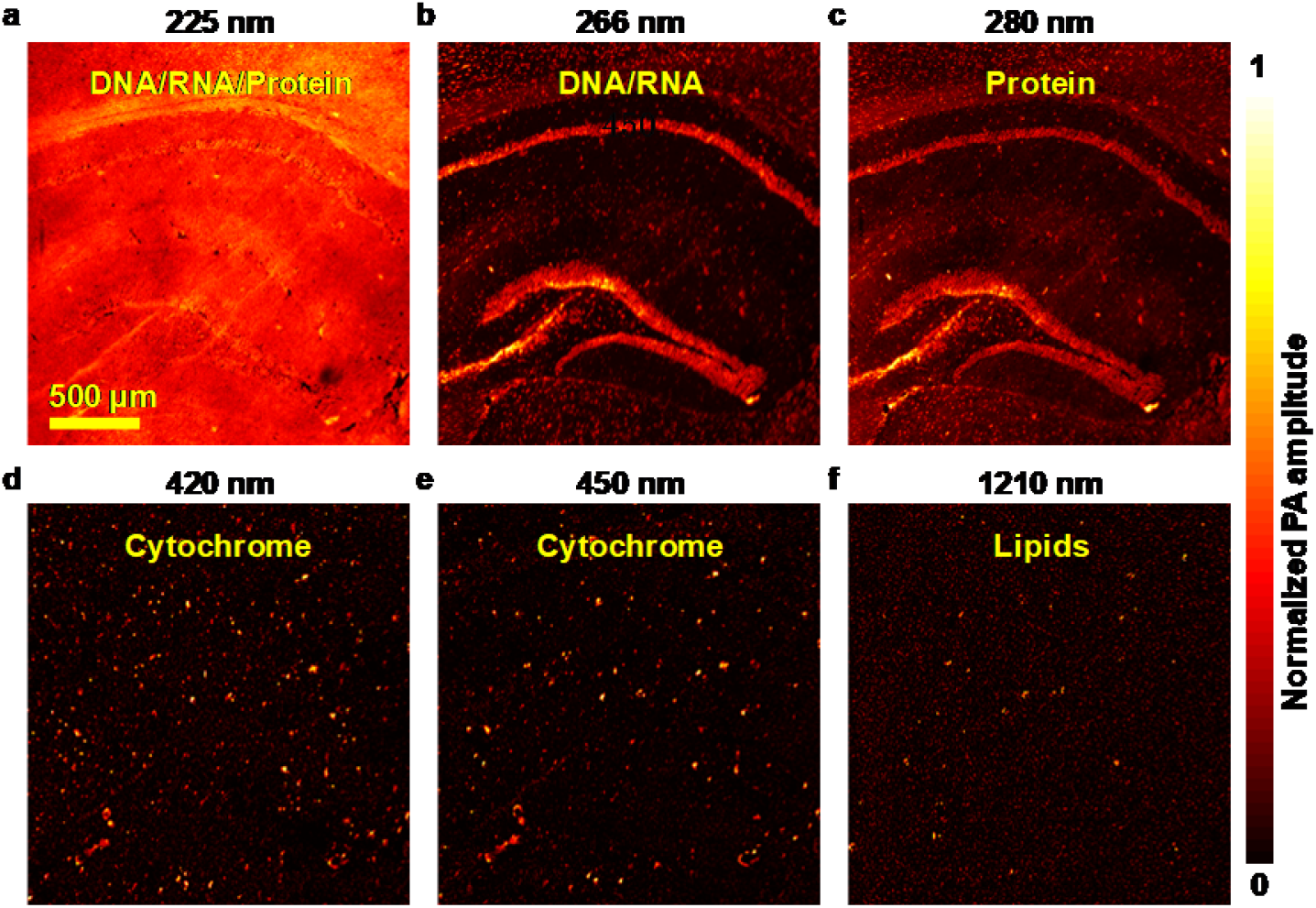
Label-free multi-wavelength wbOR-PAM images of a formalin-fixed 5-μm-thick mouse brain slice at six different wavelengths. **(a)** DNA/RNA and protein show strong absorption at 225 nm. **(b-f)** The remaining five wavelengths illustrate the distribution of DNA/RNA, protein, cytochrome, and lipids at 266, 280, 420, 450, and 1210 nm wavelengths, respectively.

To further validate the performance of wbOR-PAM for efficient cellular imaging, we imaged agarose-embedded 300-μm-thick mouse brain tissue with wavelengths ranging from 225 to 1800 nm. To image the whole brain, the region of interest was set to 6.2 mm × 6.2 mm with a lateral scanning step size of 1.6 μm. Characteristic features of cell nuclei, Nissl body, and protein were acquired at different wavelengths (Fig. 3). A wbOR-PAM image of the unstained mouse brain slice was acquired at 1720 nm, which clearly shows the structure of the brain in the coronal view (Fig. S1). As reported recently, the first and second overtones of C–H bonds of lipids are located at 1720 and 1210 nm, respectively. Excitation at these two wavelengths was utilized to image tissue abundant in lipids. Two high-contrast images of the 300-μm-thick mouse brain tissue were obtained at these two wavelengths, respectively (Fig. 3a).

**Fig. 3.**
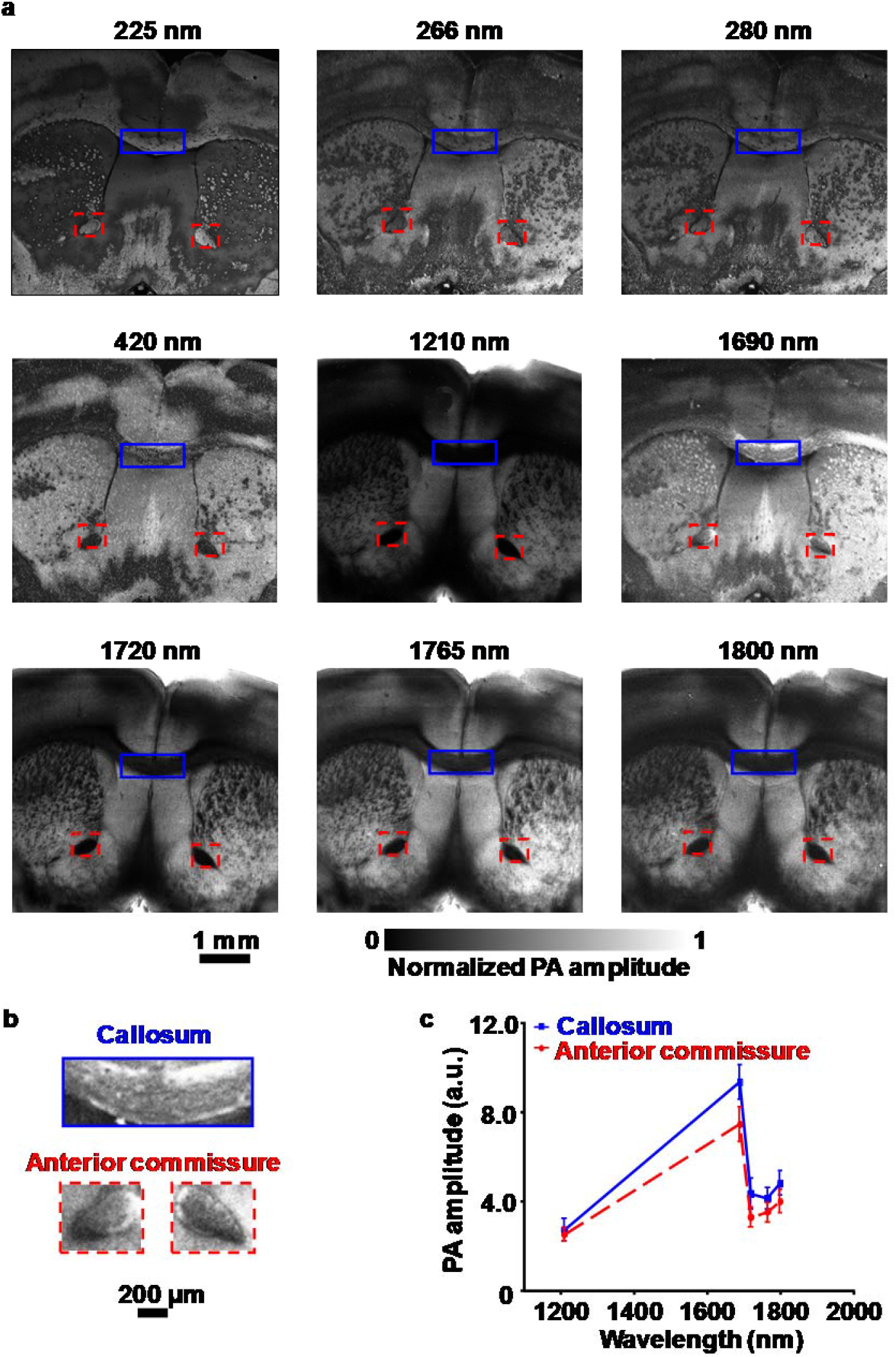
**(a)** wbOR-PAM images of a formalin-fixed 300-μm-thick mouse brain slice acquired with wavelengths of 225–1800 nm. **(b)** Zoomed-in wbOR-PAM images of callosum and anterior commissure shown in **(a)** at 1690 nm. **(c)** PA amplitude against wavelength plot of the callosum and anterior commissure by computing the averaged values of the features from **(a)**.

Phospholipids, triglycerides, and cholesterol, have been recognized to exhibit similar absorption spectra in the visible region but distinctive characteristic absorption peaks in the SWIR region. Due to the spectral similarity of chemical components in the overtone vibration region, it is challenging to identify molecules using PA signals at a single wavelength. To acquire more identifiable features, we use wbOR-PAM to map cellular chromophores with multiple laser wavelengths. As shown in Fig. 3a, distinctive contrast was acquired at 1690 nm. Close-up label-free wbOR-PAM images of callosum and anterior commissure are shown in Fig. 3b. wbOR-PAM imaging of brain slices was repeated thrice and exhibited identical images. Callosum and anterior commissure showed stronger PA amplitude at 1690 nm relative to other wavelengths (Fig. 3c). Phospholipids are the most abundant lipids in the white matter of the brain, and both callosum and anterior commissure are white matter. Furthermore, to confirm the spectral profile of endogenous contrasts, the absorption spectra of myelin and phosphatidylethanolamine were recorded from 1000 to 1800 nm (Fig. S2). Blue dashed lines in the spectra denote the absorption coefficients of these endogenous contrasts at 1690 nm. Myelin, which forms the myelin sheath of the axon in the nervous system^32^, exhibited strong absorption at 1690 nm. Compared with other absorbers (collagen^33^, human fat^34^, triglyceride^35^, and cholesterol^33^), only phospholipids exhibit strong absorption at this particular wavelength, which further highlights the capability of wbOR-PAM for imaging phospholipids without labeling.

Histopathological examination was performed to analyze the distribution of endogenous molecules in mouse brains. The stained images were acquired to compare with label-free wbOR-PAM images acquired at different wavelengths. As alcohol removes most fats for Oil Red O staining, frozen sections were prepared to examine the distribution of fats. Fig. S3 shows the wide-field microscopic images of stained frozen sections. The location of myelin was similar to that of lipids (Fig. S3c and d), which further demonstrated the low specificity of histopathological staining in differentiating phospholipids from other subsets of lipids. Furthermore, the formation of ice crystals may damage the cellular structure; thus, limitations of frozen sections including cell deformation and inaccurate localization must be taken into consideration. Fig. 4 shows the myelin in formalin-fixed slices in the wbOR-PAM image, which matched well with the stained image of the paraffin-embedded slices. Due to the prominent characteristic light absorption, phospholipids could be distinctively imaged with the wbOR-PAM system. Collectively, these results confirmed that the wavelength of 1690 nm was highly suitable for PA imaging of phospholipids without exogenous labeling and staining.

**Fig. 4.**
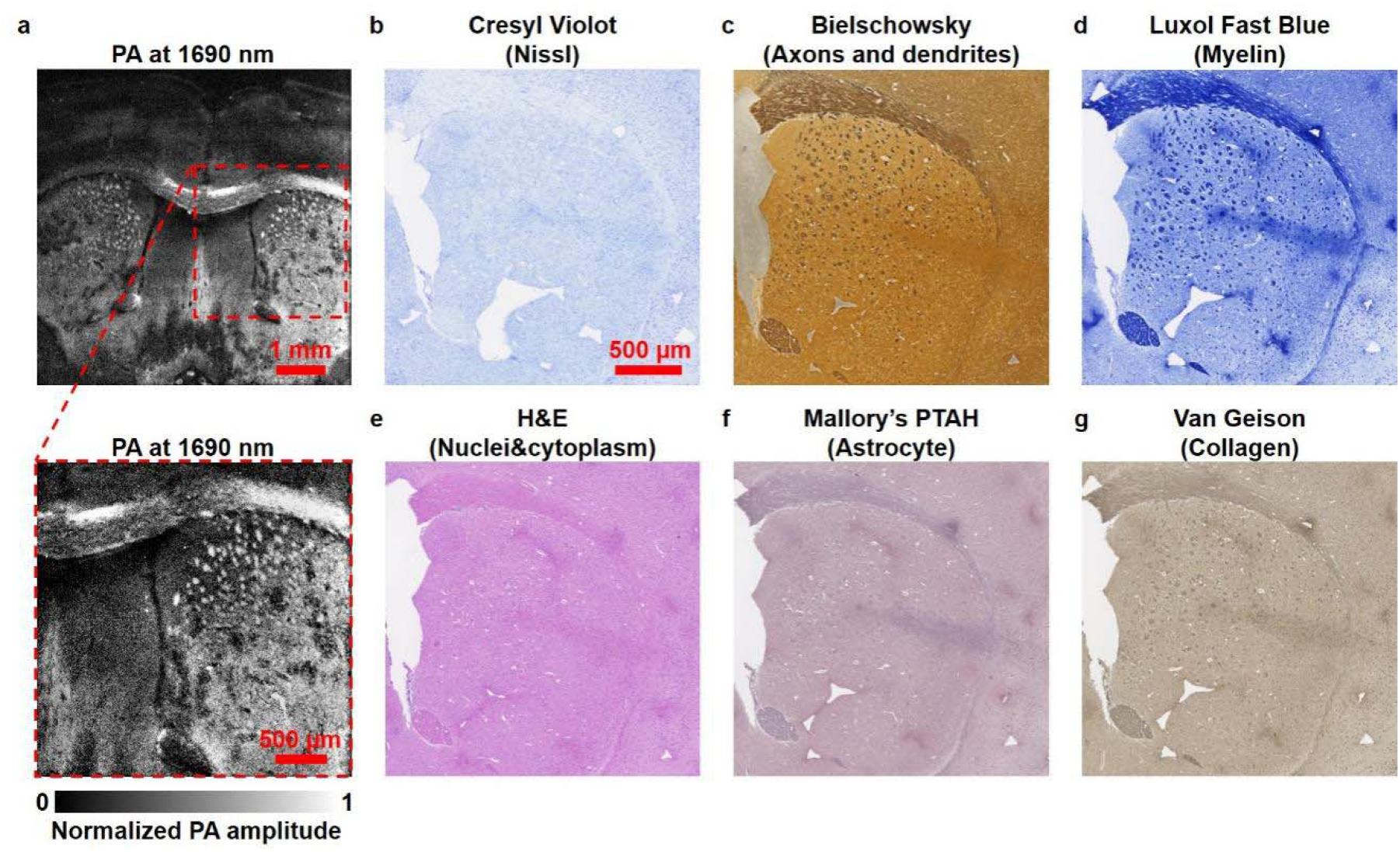
Comparison of mouse brain slices imaged by wbOR-PAM and conventional staining methods. **(a)** Label-free wbOR-PAM image of a formalin-fixed mouse brain slice acquired at 1690 nm. The image below **(a)** shows a close-up view. Histological images of paraffin-embedded brain slices stained with different dyes to highlight **(b)** Nissl, **(c)** Axons and dendrites, **(d)** Myelin, **(e)** Cell nuclei and cytoplasm, **(f)** Astrocyte, and **(g)** Collagen. **(b)–(g)** share the same scale bar (500 μm).

The cell nuclei, phospholipids, and lipids in the wbOR-PAM images were further processed and highlighted in green, red, and blue, respectively (Fig. 5). Fig. S4a and b illustrate wbOR-PAM images obtained at 266 nm and 1690 nm, which sketched the outline of cell nuclei and phospholipids of a 300 μm-thick formalin-fixed mouse brain slice. To show the distributions of cell nuclei and phospholipids in a single image, the two images were overlaid (Fig. S4c). The corresponding close-up images of cell nuclei and phospholipids are shown in Fig. S4d. The distance and the diameter of phospholipids were calculated for wbOR-PAM images. These results demonstrated that the distribution and the calculated diameter of phospholipids were different from that of cell nuclei (Fig. S5). The wbOR-PAM images of consecutive coronal slices were digitally extracted through the 300 μm-thick brain slices with high resolution and deep imaging depth (118 μm). In contrast with imaging a thin brain slice, imaging an unstained thick specimen resulted in revealing multilayered information without physical sectioning (Fig. S6). Therefore, by using sets of wavelengths, the designed wbOR-PAM system can be applied as an effective tool to reveal additional crucial biomolecules and provide better foundations to study the metabolism of life.

**Fig. 5.**
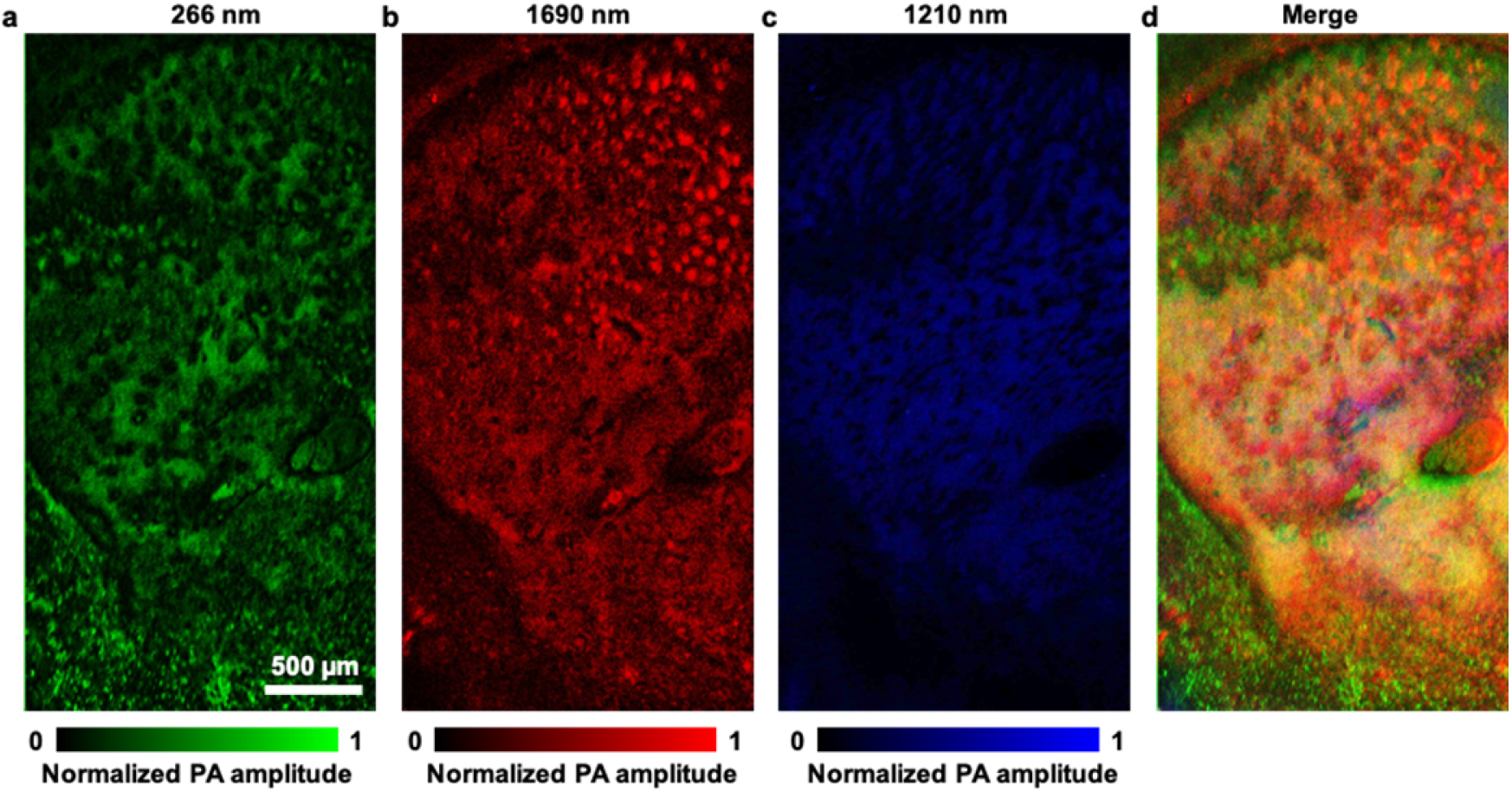
wbOR-PAM images of cell nuclei, phospholipids, and proteins in a 300 μm-thick formalin-fixed mouse brain slice. Label-free wbOR-PAM image obtained at **(a)** 266 nm (green), showing cell nuclei; **(b)** 1690 nm (red), illustrating the location of phospholipids; **(c)** 1210 nm (blue), displaying the distribution of proteins. **(d)** Merged images of **(a)–(c)**.

## Conclusion and Discussion

wbOR-PAM comprehensively analyzes cellular components with tunable wide-band wavelengths ranging from 210 to 2600 nm. In wbOR-PAM, we used a reflective objective lens that is free of chromatic aberration, which enables stable and continuous confocal optical-acoustic alignment over the entire UV to SWIR regions. Compared with existing OR-PAM systems, wbOR-PAM empowers detailed pathological and structural analysis owing to the various choices of wavelengths for specific absorption bands of biomolecules.

In this study, we imaged mouse brain tissue using nine wavelengths ranging from the UV to the SWIR regions (225, 266, 280, 420, 1210, 1690, 1720, 1765 and 1800 nm). These wavelengths were selected to differentially highlight the contrasts among DNA/RNA, protein, cytochrome, phospholipids and lipids. Merging wbOR-PAM images does not require digital registration since no sample processing or staining is needed. With the addition of specific wavelengths, wbOR-PAM exhibit potential to capture additional major biomolecules and cellular components, including phospholipids at its characteristic absorption wavelength as demonstrated here. Phospholipids account for the highest proportion of lipids in the brain and have been significantly associated with brain diseases such as demyelinating disease^36^, atherosclerotic lesions^37^, and sphingomyelin lipidosis^38^. Thus to efficiently capture the alterations of phospholipids, it is essential to design sensitive phospholipid-fingerprinting methods. Herein, using wbOR-PAM, the molecular components of complex brain slices were probed. The findings of the present study demonstrated a distinctive absorption of phospholipids at 1690 nm and revealed their distribution in wbOR-PAM images, for the first time. Notably, myelin-stained images also verified the results of wbOR-PAM with high consistency. Thus, wbOR-PAM provides a new way to analyze disease-related alterations in cellular components without labeling and to study disease pathogenesis.

## Materials and Methods

### System setup

A schematic of our wbOR-PAM system is illustrated in Fig. 1. A tunable OPO laser (NT242-SH, EKSPLA Ltd., Lithuania) is used as the light source. The OPO, pumped by a Nd:YAG (neodymium-doped yttrium aluminum garnet) laser (repetition rate = 1 kHz), covers a wide tunable wavelength range from 210 to 2600 nm with a pulse duration of 3–10 ns. The beam profile of the laser beam is improved by a spatial filter that is composed of a pair of parabolic mirrors and a 10-μm-diameter pinhole. Subsequently, the laser beam is illuminated on the specimen through a reflective objective lens (#59-886, Edmund Optics, Inc. USA), which is confocally-aligned with the focal zone of an ultrasonic transducer in transmission mode. PA signals are recorded by the ultrasonic transducer with a center frequency of 50 MHz (V214-BC-RM, Olympus NDT, Inc., Japan). Thus, an optical diffraction-limited lateral resolution that could be achieved with the wbOR-PAM system was defined as *λ*/2NA, where *λ* is the wavelength used to image the specimen and NA is the numerical aperture of the objective lens. The axial resolution is about 30 μm, which is determined by the bandwidth of the ultrasonic transducer. Wavelength tuning is set by a control pad. PA data are obtained by a high-speed data acquisition card (ATS9350, Alazar Technologies Inc., Canada). The motor movement, laser triggers, and data acquisitions are synchronized using a customized LabVIEW program.

### Mouse brain tissue preparation

All the animal experiments procedures were approved by both the Animal Studies Committees of California Institute of Technology, USA. Mouse brains were collected from euthanized Swiss Webster mice (Hsd: ND4, Harlan Laboratories, Indiana). Post-harvest, brain tissue specimens were immediately fixed with formalin and submerged in agarose to maintain its shape. The fixed brain specimens were coronally sectioned into thin slices with different sectioning thicknesses on a vibrating blade microtome (VT1200 S, Leica Biosystems Inc., Germany). After PA imaging, the corresponding histologic sections were sent for histologic examination. Serial sectioning method was performed for a collection of three-dimensional microstructural data^39^. Multiple staining techniques were implemented for different slices of the same specimen to prevent interference due to different staining methods.

### Spectral analysis

For spectral analysis, myelin and phosphatidylethanolamine were obtained from Sigma-Aldrich, Inc. UV/VIS/NIR spectrophotometry (Lambda 950, PerkinElmer Inc., USA) with a wavelength range of 175– 3300 nm was used to determine the absorption spectra. To capture various endogenous and differentiate diverse lipid classes, PA signals were collected from different mouse brain slices at different wavelengths. For optical spectral analysis, a DWARF-Star spectrometer (StellarNet, Inc., USA) was used to calibrate the wavelength of the OPO laser. The parameter was quantified with the SpectraWiz^®^ software.

### wbOR-PAM of fixed brain slices

For PA imaging, a UV laser was selected to analyze DNA/RNA^40^. The unstained brain specimens were placed on UV transparent quartz slides (19 mm × 19 mm, Chemglass, Inc, USA) instead of glass slides. For PA imaging of lipids, heavy water was added in a water tank for acoustic coupling, as regular water has strong absorption in the NIR region. After sectioning, the specimen on a quartz slide was sandwiched between two plastic membranes of the water tank and placed on a laboratory-made specimen holder. The laser beam was focused on the bottom surface of the specimen, and PA signals were detected by a confocally-aligned ultrasonic transducer.

### Histologic staining and imaging

After wbOR-PAM imaging, histologic staining was performed to highlight specific tissue components of the brain slices. Stained sections were prepared by Histoserv, Inc (USA). The formalin-fixed brain specimens were embedded in paraffin and sectioned. Eight serial thin slices from the same block were processed with different staining protocols. H&E staining was used to reveal cell morphology. Myelin was stained with Luxol Fast Blue stain. The Bielschowsky silver impregnation method was applied to image axons and dendrites. Nissl substances were evaluated by the Cresyl Violet staining method. Astrocytes were stained with phosphotungstic acid-hematoxylin. The Von Kossa and Warthin-Starry methods were used for histological visualization of calcium and melanin, respectively. As lipids are soluble in organic solvents, the standard paraffin staining method is not suitable for Oil Red O staining. The thick brain slice was embedded in an optimum cutting temperature compound for cryosectioning. The frozen sections were stained with Oil Red O. The stained sections were then viewed with a bright-field microscope (NanoZoomer, Hamamatsu Photonics K. K., Japan), which is an optical line scanning system that allows imaging the entire slide with up to 40× close-up in a minute. These stained images were compared with the wbOR-PAM images.

## Supplemental document

See Supplementary Materials for supporting content.

## Supplementary Information

**Fig. S1.**
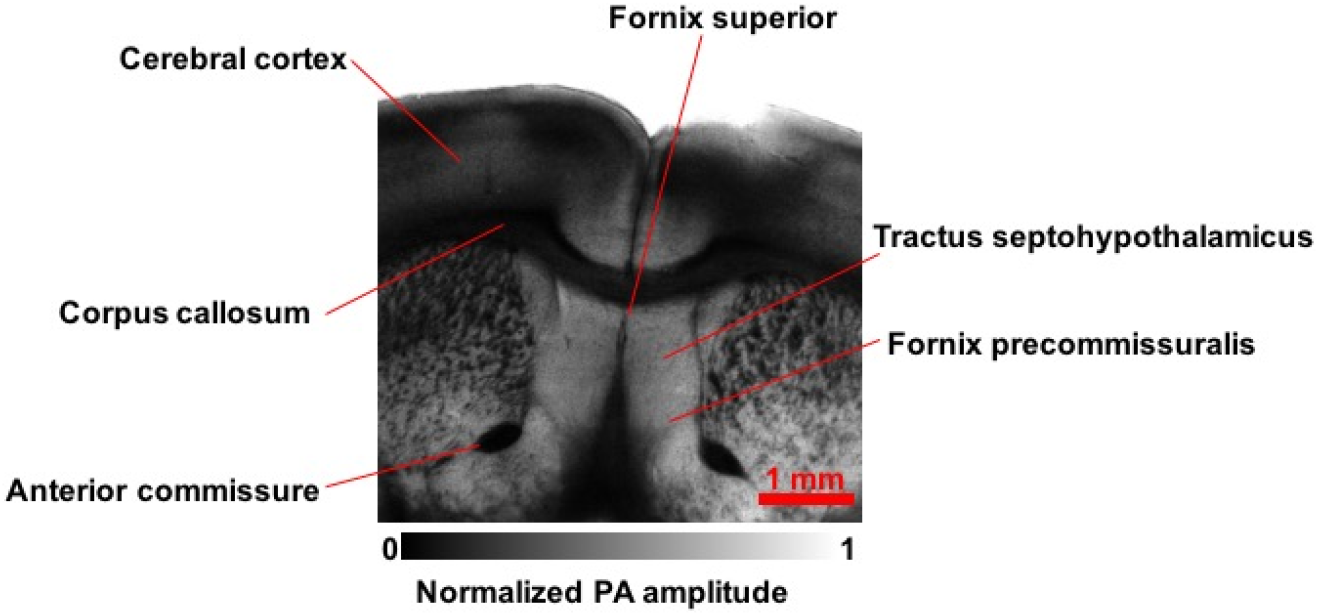
Label-free wbOR-PAM image of a mouse brain slice acquired at 1720 nm, showing the structure of a mouse brain in coronal view clearly.

**Fig. S2.**
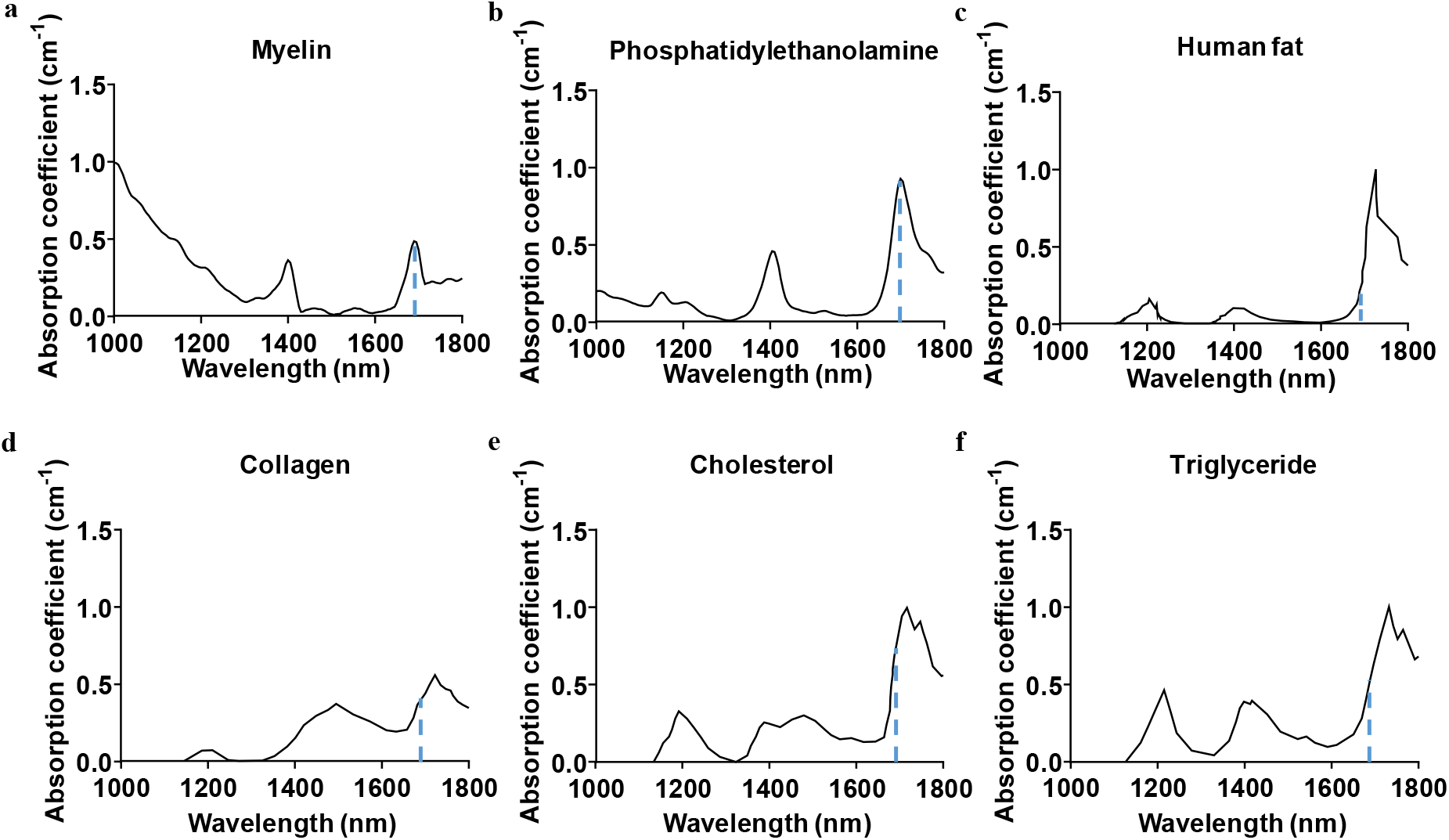
Absorption spectra of **(a)** myelin, **(b)** phosphatidylethanolamine, **(c)** human fat, **(d)** collagen, **(e)** cholesterol, and **(f)** triglyceride. Blue dashed lines in the spectra denote the absorption coefficients of endogenous contrasts at 1690 nm.

**Fig. S3.**
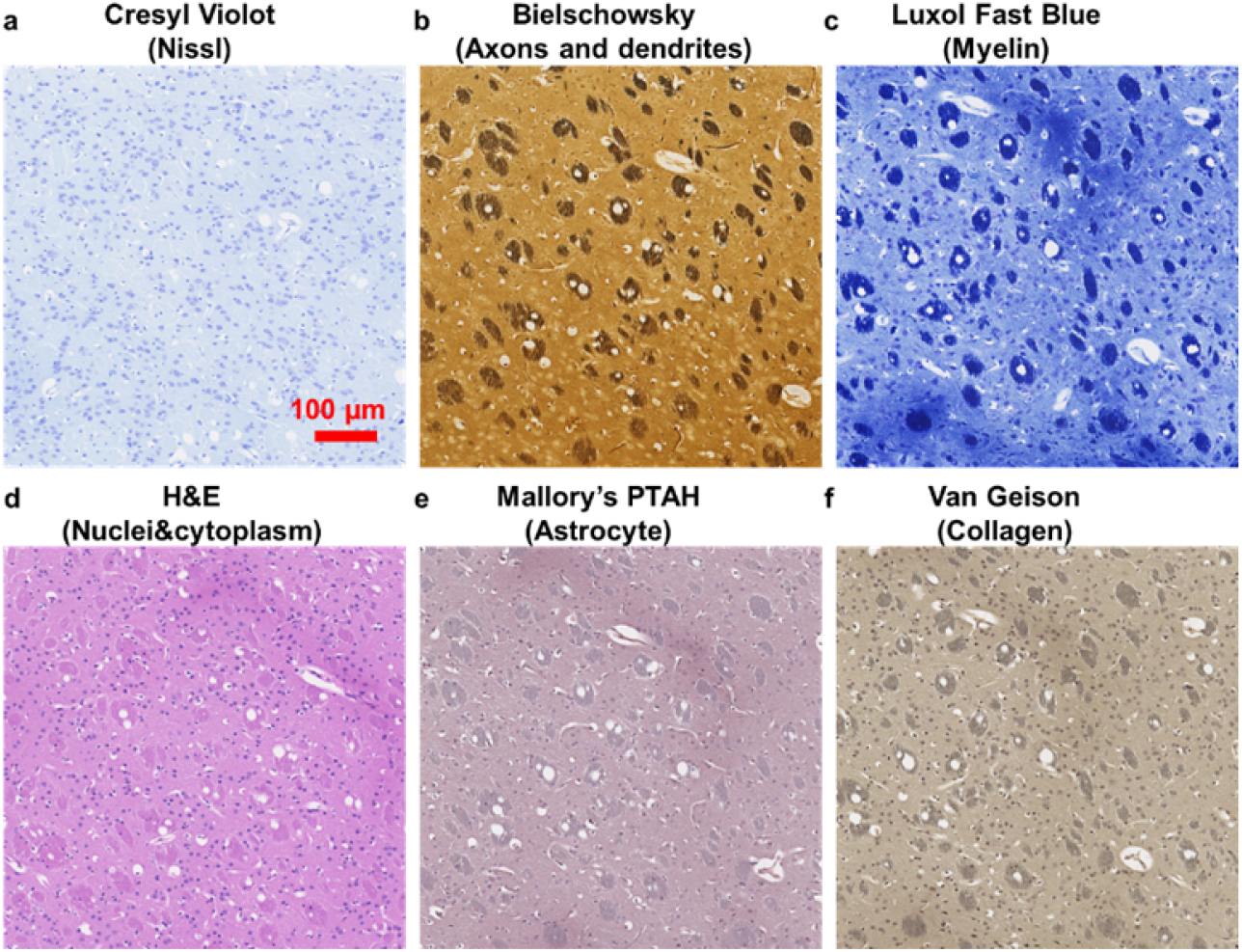
Histological images of paraffin-embedded brain slices stained with different dyes to highlight **(a)** Nissl, **(b)** Axons and dendrites, **(c)** Myelin, **(d)** Cell nuclei and cytoplasm, **(e)** Astrocyte, and **(f)** Collagen. **(a)–(f)** share the same scale bar (500 μm).

**Fig. S4.**
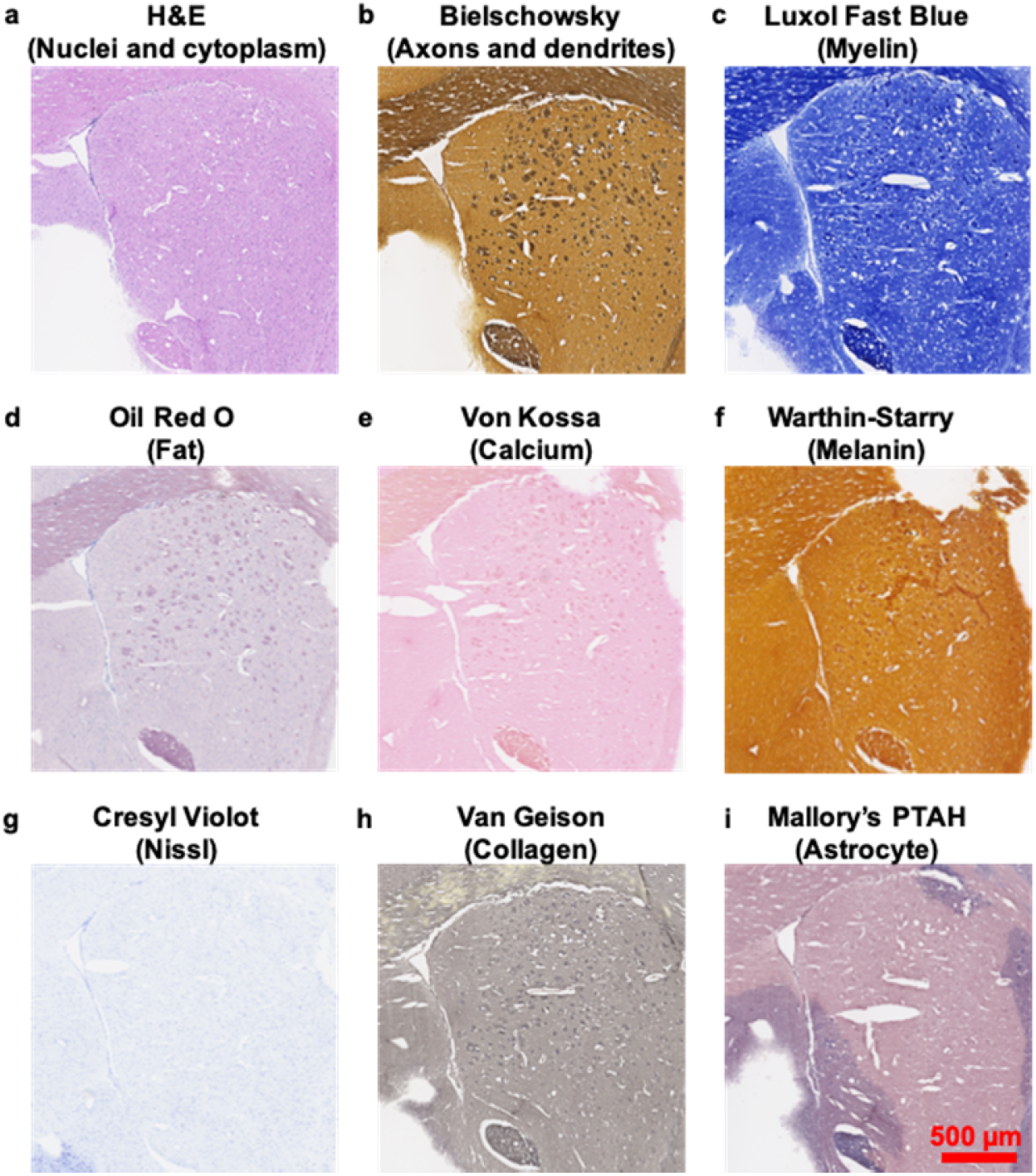
Histological images of 10-μm thick frozen brain slices stained with different dyes showing **(a)** Nuclei and cytoplasm, **(b)** Axons and dendrites, **(c)** Myelin, **(d)** Fat, **(e)** Calcium, **(f)** Melanin, **(g)** Nissl, **(h)** Calcium, and **(i)** Astrocyte. All the images share the same scale bar marked in **(i)**.

**Fig. S5.**
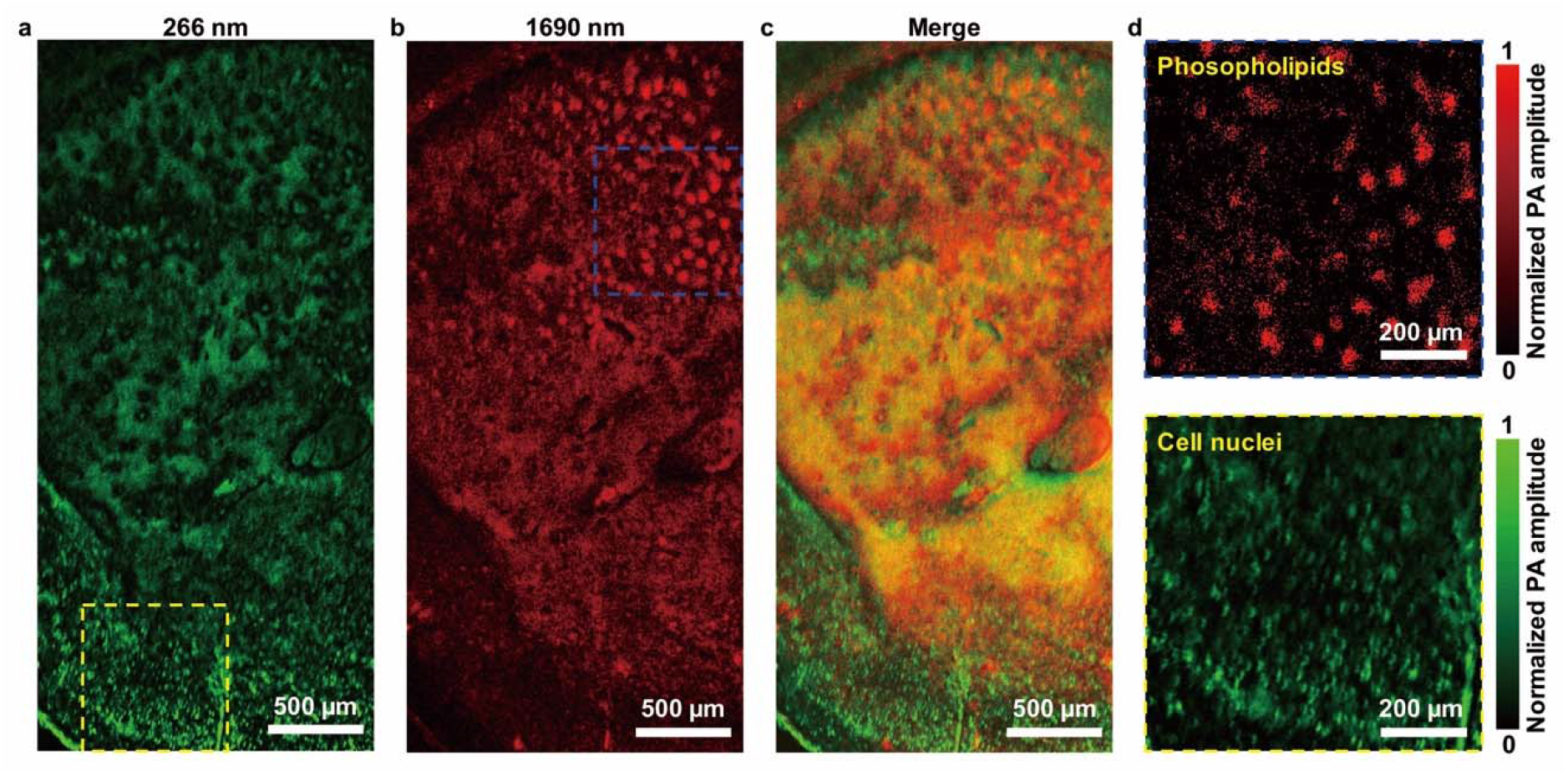
Cell nuclei and phospholipids in wbOR-PAM images of a 300 μm-thick formalin-fixed mouse brain slice. **(a)** wbOR-PAM image obtained at 266 nm (green) shows cell nuclei. **(b)** wbOR-PAM image obtained at 1690 nm (red) shows the distribution of phospholipids. **(c)** Merged images of **(a)** and **(b). (d)** Close-up images of phospholipids (blue dashed square) in **(b)** and cell nuclei (yellow dashed square) in **(a)**, respectively.

**Fig. S6.**
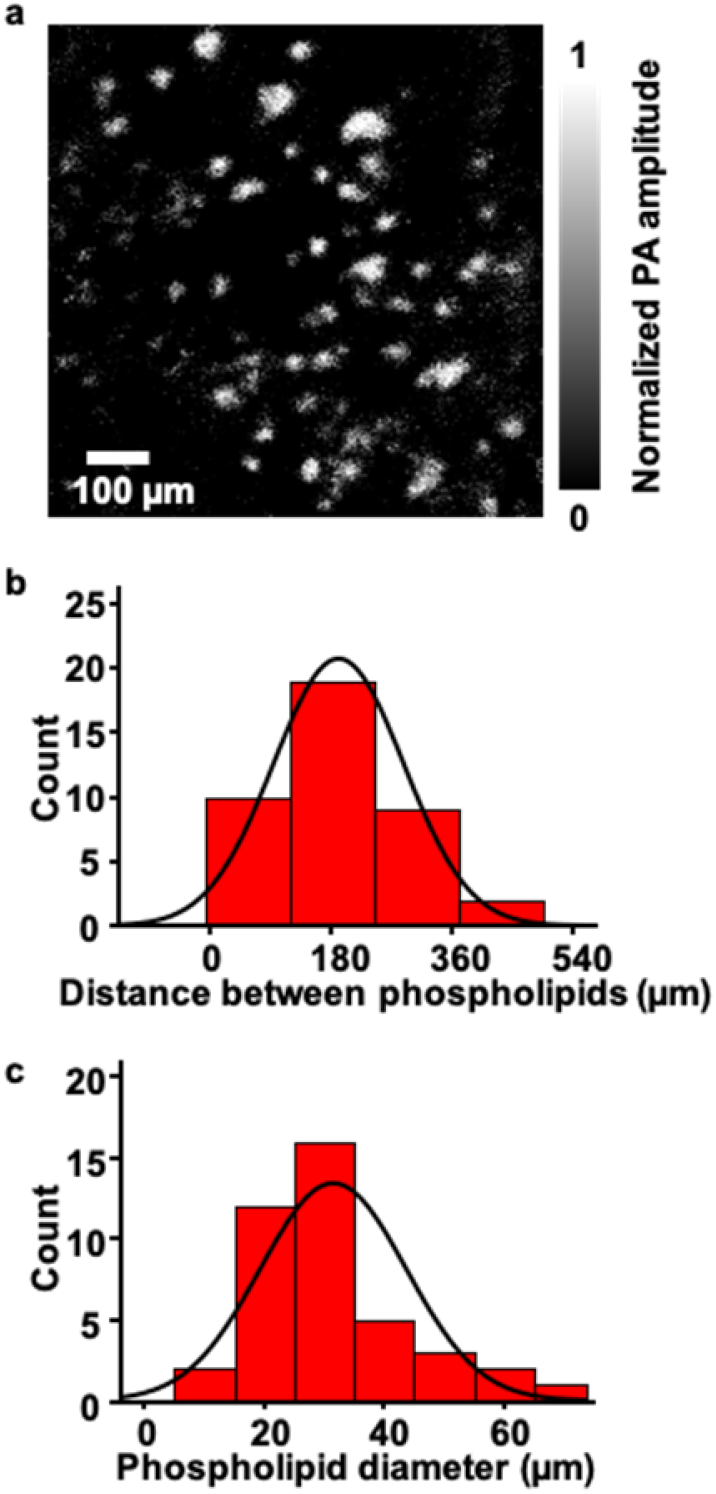
Distributions of phospholipids in the mouse brain slice. **(a)** wbOR-PAM image of phospholipids, obtained at 1690 nm. **(b)** Histogram of the distance between phospholipids (**n**= 40). The black curve is a Gaussian fit with a mean of 190.7 μm and a standard deviation (SD) of 96.6 μm. **(c)** Histogram of the phospholipids diameter (**n**= 41). The black curve is a Gaussian fit with a mean of 31.6 μm and an SD of 12.2 μm.

**Fig. S7.**
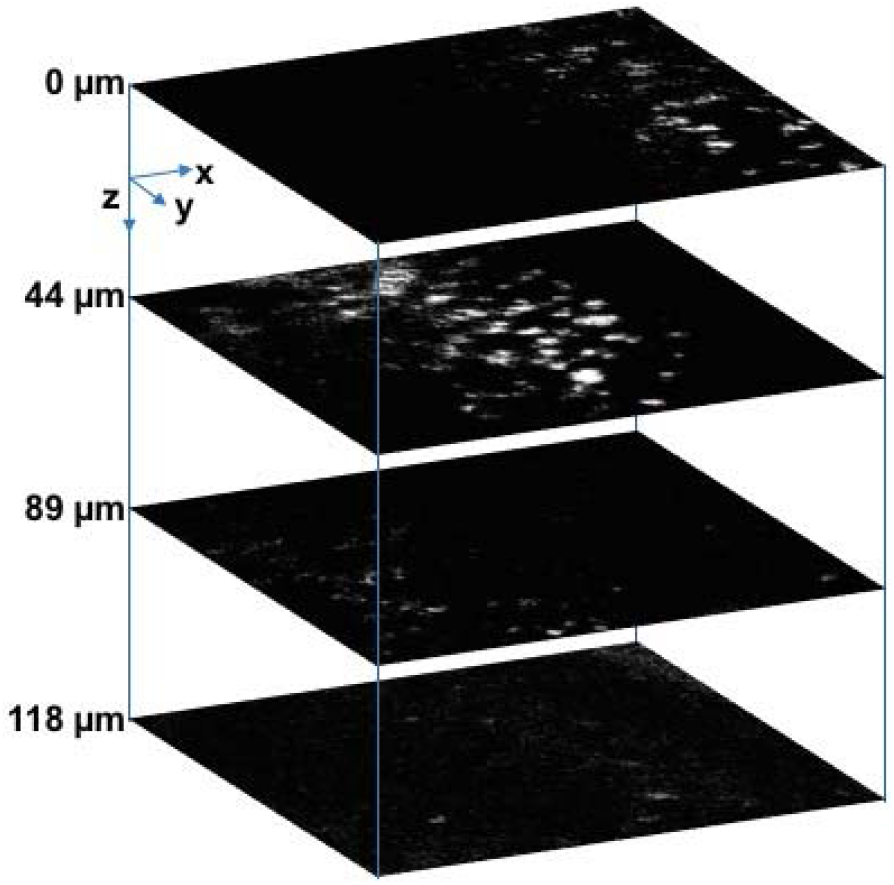
Label-free wbOR-PAM images of phospholipids in a 300 μm-thick mouse brain slice acquired at 1690 nm in different layers (depths of 0, 44, 89 and 118 μm).

